# Bacteria in Paper, a versatile platform to study bacterial ecology

**DOI:** 10.1101/464347

**Authors:** Felix JH Hol, George M Whitesides, Cees Dekker

## Abstract

Habitat spatial structure has a profound influence on bacterial life, yet there currently are no low-cost equipment-free laboratory techniques to reproduce the intricate structure of natural bacterial habitats. Here, we demonstrate the use of paper scaffolds to create landscapes spatially structured at the scales relevant to bacterial ecology. In paper scaffolds, planktonic bacteria migrate through liquid filled pores, while the paper’s cellulose fibers serve as anchor points for sessile colonies (biofilms). Using this novel approach we explore bacterial colonization dynamics in different landscape topographies, and characterize the community composition of *Escherichia coli* strains undergoing centimeter-scale range expansions in habitats structured at the micrometer scale. The bacteria-in-paper platform enables quantitative assessment of bacterial community dynamics in complex environments using everyday materials.

---

The intricate spatial structure of microbial habitats has a decisive influence on the populations they support. Many habitats, including biological tissues and soil matrices, consist of a microscale network of connected pores and cavities through which cells migrate, while their abundant surfaces facilitate the growth of biofilms. Physical and chemical heterogeneities through space give rise to the diverse and architecturally complex microbial communities we find in nature [1–6]. The smallest ecological scale at which microbes interact with their environment is set by the size of the organisms and is on the order of micrometers, while environmental gradients extend over millimeters and beyond. Traditional laboratory tools to culture microbes are not well suited to mimic realistic landscapes at those scales, and furthermore typically only support planktonic *or* surface-associated growth (not both simultaneously), and thus suppress the coexistence of these distinct lifestyles. In the past decade, various microfabrication-based approaches to culture microbes have emerged, enabling the study of bacterial ecology at the micrometer to millimeter scale by engineering synthetic landscapes. While such approaches have resulted in exciting insights regarding e.g. spatial competition between microbes, the evolution of antibiotic resistance, microbial community assembly, and biofilm growth [3, 7–18], microfabrication-based approaches to study bacterial ecology have not been adopted widely. This is largely due to the fact that the laboratory infrastructure necessary to create microfabricated landscapes is expensive, specialized, and not readily available in microbiology labs.

To overcome this barrier, we here demonstrate the use of simple paper scaffolds as a versatile and easy-to-use platform for studying bacterial communities in environments that are spatially structured at the relevant microscopic scales. Paper is a widely available material consisting of cellulose fibers. Interestingly, the characteristic length scales of paper [19] and many bacterial habitats (e.g. the soil matrix [20]) are very similar, having pores from a few to several tens of micrometers. Furthermore, paper can be easily cut, either by hand or using a laser cutter, into any two-dimensional geometry at the millito centimeter scale, while layers of patterned paper can be stacked to make three-dimensional geometries [21–24]. Paper furthermore can be creased and folded to create yet other geometries. The spatial scales at which paper is structured (microns) and can be manipulated (millito centimeter) correspond very well to the range of scales that are intrinsic to bacterial ecology, suggesting that paper may provide an excellent substrate for mimicking the complex structure of natural bacterial habitats.

## RESULTS & DISCUSSION

To facilitate bacterial growth and motility in a paper matrix, we saturated paper with bacterial growth medium (LB broth). Confocal imaging of fluorescently labelled *Escherichia coli* demonstrated that bacteria can swim in the liquid medium that fills pores in the cellulose mesh, allowing bacterial growth, migration, and colonization throughout paper scaffolds several centimeters in length. Figure 1 shows sessile colonies (biofilms) formed by *E. coli* (Fig. 1B,C) and *Bacillus subtilis* (Fig. 1D) after a 15 hour incubation period at 37°C. Cellulose fibers act as anchor points for surface associated growth, giving rise to dense colonies that form in the pores (see Supplementary Movie 1 for a confocal Z-stack showing bacterial aggregates that formed 0–33 *µ*m into the paper). Before inoculation of bacteria at one central point of the paper scaffold, the entire scaffold was saturated with growth medium forming an initially homogeneous nutrient landscape. Yet strong demographic heterogeneities were apparent after 15 hours of growth when cellular aggregates had formed scattered throughout the paper scaffold – even at the extremities, centimeters away from the original inoculation point.

**FIG. 1.**
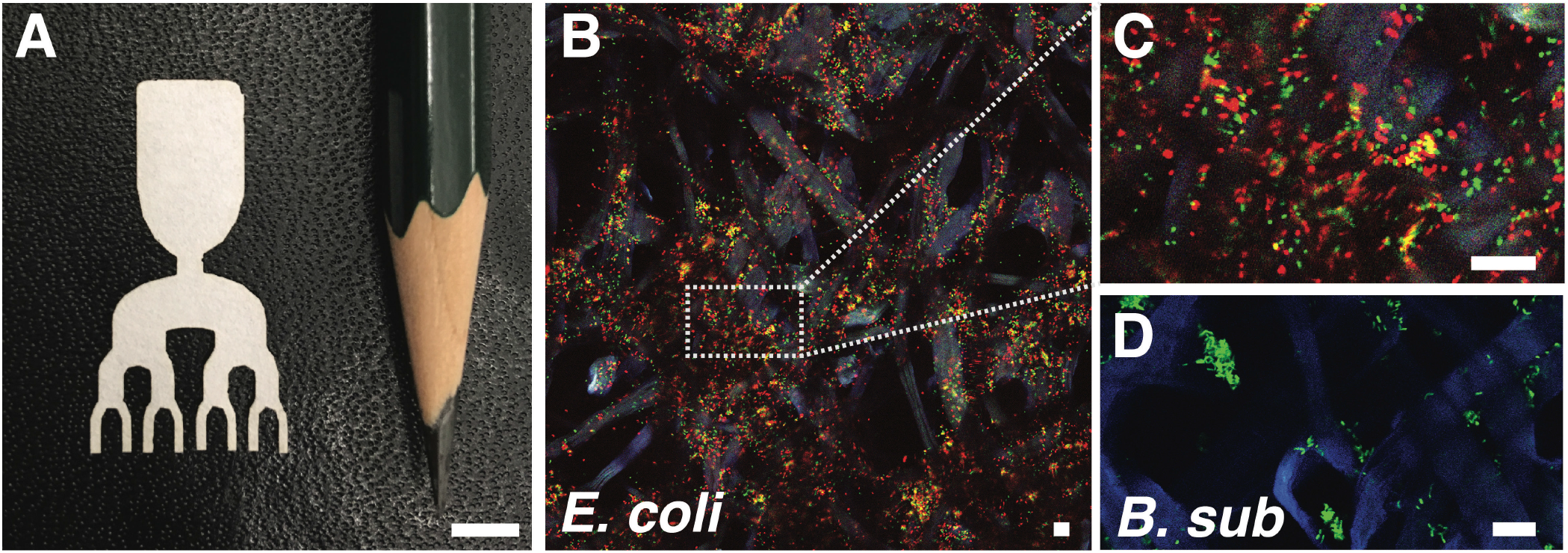
Bacteria-in-paper. A) A photograph showing a paper scaffold cut to a predesigned shape with a laser cutter. A pencil is shown for scale, the scale bar is 5 mm. B) Confocal scan of bacteria in paper showing *GFP* expressing *E. coli* (green), *RFP* expressing *E. coli* (red), and paper (blue). C) Zoom in of the area indicated with dashed lines in (C). D) Confocal scan of *GFP* expressing *B. subtilis* (green) and paper (blue). Scale bars in B-D are 20 *µ*m.

Confocal imaging penetrates up to *∼*100 micrometers into the paper, enabling the high-resolution visualization of communities of fluorescently labelled bacteria inhabiting the paper. The paper in use here weighs 87 g/m^2^ and is 180 micrometers thick, images taken at multiple focal distances (Z-stacks) from both sides can thus be used to visualize the entire community. However, cellulose fibers may obscure a fraction of cells when imaging beyond several tens of micrometers into the paper. To enable quantitative assessment of the bacterial communities inhabiting the paper matrix independent of the penetration depth of imaging, we took advantage of the fact that bacterial DNA can easily be extracted from paper to assess the community composition by e.g. quantitative PCR (qPCR) or sequencing-based methods (e.g. [25]). As we demonstrate below, qPCR provides an economical and convenient means to spatially resolve community composition, albeit at a lower resolution compared to confocal microscopy.

Having established that bacteria are motile and grow in paper containing growth medium, we used this approach to investigate the colonization dynamics of *E. coil* in a range expansion in two distinct types of landscapes. The connectivity, or network topology, of a landscape is known to influence community dynamics and biodiversity across all taxa [26–28]. Branching networks, such as rivers or cave systems, comprise a ubiquitous class of landscapes that can also be found at microscopic scales in e.g. lungs, capillary networks, or soil. To probe the effect of a branching landscape topology on bacterial range expansions, we cut paper scaffolds (26 x 15 mm) that consist of a central inoculation zone providing access to both a branching, and a non-branching landscape on opposite sides (top and bottom in Figure 2A, respectively). As both landscapes are colonized from the same inoculation zone, and thus by the same initial community, the effect of branching on the range expansions can be assessed by comparing the community composition at the extremities of both landscapes. Paper scaffolds were saturated with rich growth medium (LB) and inoculated in the center with a 1:1 mixture of neutrally labelled *E. coli*, isogenic except for a green fluorescent protein (GFP) versus red fluorescent protein (RFP) insertion in the *Lac* operon [3, 10, 29]. A challenge to confining bacteria to liquid-saturated paper is the liquid film that forms when wet paper comes in contact with a surface (e.g. a glass coverslip). In order to prevent bacteria from growing in or migrating through such a liquid film, we suspended the wet paper on thin wires (*∼*5 mm pitch) in a chamber with saturated humidity. This ensures no liquid interfaces are formed and all bacterial migration happens through the paper matrix.

Confocal imaging of the branches demonstrated that bacteria successfully colonized the full length of both landscapes during a 15 hour incubation period, and revealed mixed (both colors) cellular assemblages at the branch extremities indicating coexistence of the two strains (Figure 2). To determine the community composition at the ends of the range expansion, we extracted genomic DNA from 2 mm^2^ paper fragments cut from the branch extremities. Utilizing qPCR we assessed the population fraction of GFP versus RFP labeled *E. coli* in the branches using primer pairs amplifying a fragment of the respective genes encoding for the fluorescent proteins. Quantitative PCR showed that the average (global) community composition at the branch extremities did not differ from the community composition at the far edge of the linear system (rank sum test, *p <* 0.01), nor did either average deviate from the composition of the population in the inoculation zone. Although the averages were similar, the variation in community composition between branches differed, being larger (one-sided *F*-test, *p* = 0.03) than the variation among patches of equal size in the non-branching system.

**FIG. 2.**
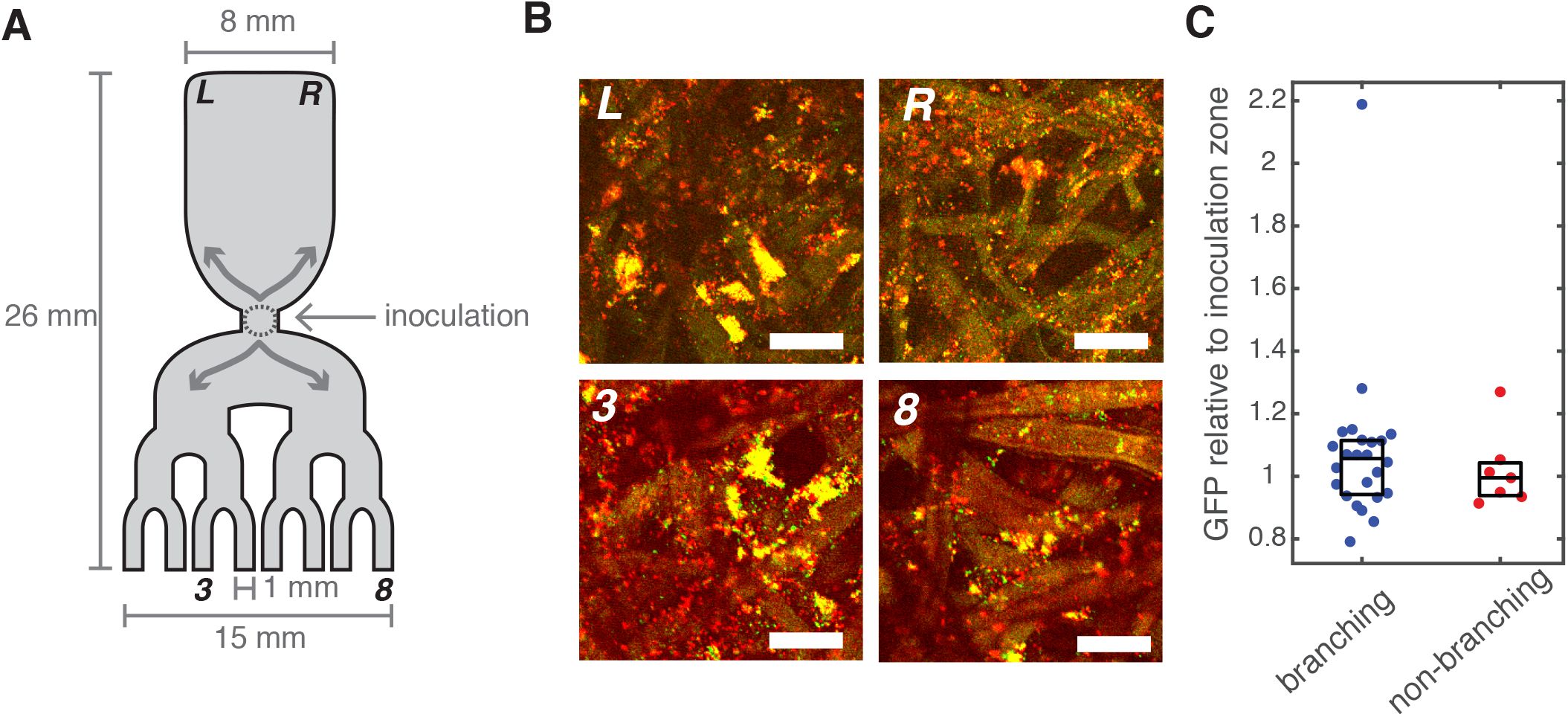
Range expansions in branching and non-branching landscapes. A) Cartoon of a paper scaffold consisting of a branching landscape and a non-branching landscape connected to the same inoculation zone (indicated by a dashed circle), arrows indicate the direction of migration and population expansion upon inoculation. B) Confocal scans of *GFP* and *RFP* labelled *E. coli* at the landscape’s extremities labelled L, R, 3, and 8 in panel A. Scale bars are 20 *µ*m. C) Fraction of *GFP* labelled *E. coli* relative to the *GFP* fraction at the inoculation zone measured at the branch extremities by qPCR on gDNA extracted from the most distal 2 mm of each branch (i.e. all 8 branches for the branching landscape, and the left- and right-most corners of the non-branching landscape). Data is plotted for 3 replicate experiments (*n* = 3), the central line indicates the median, the bottom and top edges indicate the 25th and 75th percentile, respectively.

These results are in agreement with theoretical studies suggesting that inter-branch diversity is increased in dendritic networks [26, 30]. The variation between branches we observe, however, is only moderately larger than differences between equally sized patches in the non-branching network, despite the fact that the range expansion covers centimeter distances, i.e. *∼* 10^3^ body lengths. Interestingly, the relatively balanced population composition at the branch extremities, and the low variation between branches we observe, contrasts findings from a different experimental system commonly used to study microbial range expansions, namely bacterial colonies growing on solid agar [31, 32]. *E. coli* are non-motile on solid agar, and a range expansion of two neutral strains growing on solid agar starting from a mixed point-inoculation, is governed by a stochastic coarsening process in which a small number of pioneers quickly dominates the expanding front, diminishing local diversity [31, 32]. In contrast, the current paper-based system supports local coexistence of the two strains throughout the range expansion. Two strain coexistence is even observed at micrometer scales within an individual branch (Fig. 2), suggesting that coarsening along the range expansion is completely absent in paper scaffolds. Likely, the stark differences in colonization dynamics in paper scaffolds compared to solid agar originate (in part) from the different modes of dispersal and growth that the two systems support: dispersal by growth and division only on solid agar, versus swimming motility and co-occurrence of sessile and planktonic lifestyles in paper scaffolds. It is interesting to note that the distinct lifestyles that the paper scaffolds support are an important ingredient of bacterial community assembly in natural habitats [33]. Taken together, these results suggest that when growth and division are the only modes of dispersal, this leads to a coarsening of the community composition along the range expansion, while habitats that support the full range of dispersal modes (i.e. growth and motility) promote community mixing to a much larger extent resulting in a high degree of diversity even at local scales.

We took advantage of the versatile nature of growing bacteria in paper to explore colonization in a second, quite different ecological scenario, an archipelago of islands. We constructed a landscape consisting of a ‘main-land’ (used to inoculate the system) and several uninhabited islands situated 5–15 millimeters from the mainland. The (non-inoculated) islands were impregnated with 1 microgram of glucose and dried. The mainland was inoculated with a 1:1 mix of GFP and RFP labelled *E. coli*, the archipelago was subsequently sandwiched between glass slides, and the remaining space (the ‘sea’) filled with minimal medium (lacking a carbon source). The bacteria thus initially faced a low-nutrient environment dotted with nutrient-rich islands. Upon wetting the landscape, the solid glucose slowly dissolved and diffused out of the non-inoculated islands, creating a dynamic and heterogeneous resource landscape. Figure 3 shows that after 15 hours of incubation, *E. coli* from the mainland had successfully colonized the non-inoculated paper scaffolds and established colonies in the islands. The mainland was colored yellow due to a uniform mix of green and red cells. Interestingly, a very different distribution of single-colored colonies can be seen scattered across the three islands. The single-color colonies likely derive from individual colonizers, which gave rise to distinct founder populations. Community structure at the islands thus differs from the mainland, exhibiting a much lower local diversity (i.e. the characteristic length scale of clonal single-color patches is much larger) due to relatively rare colonization events.

**FIG. 3.**
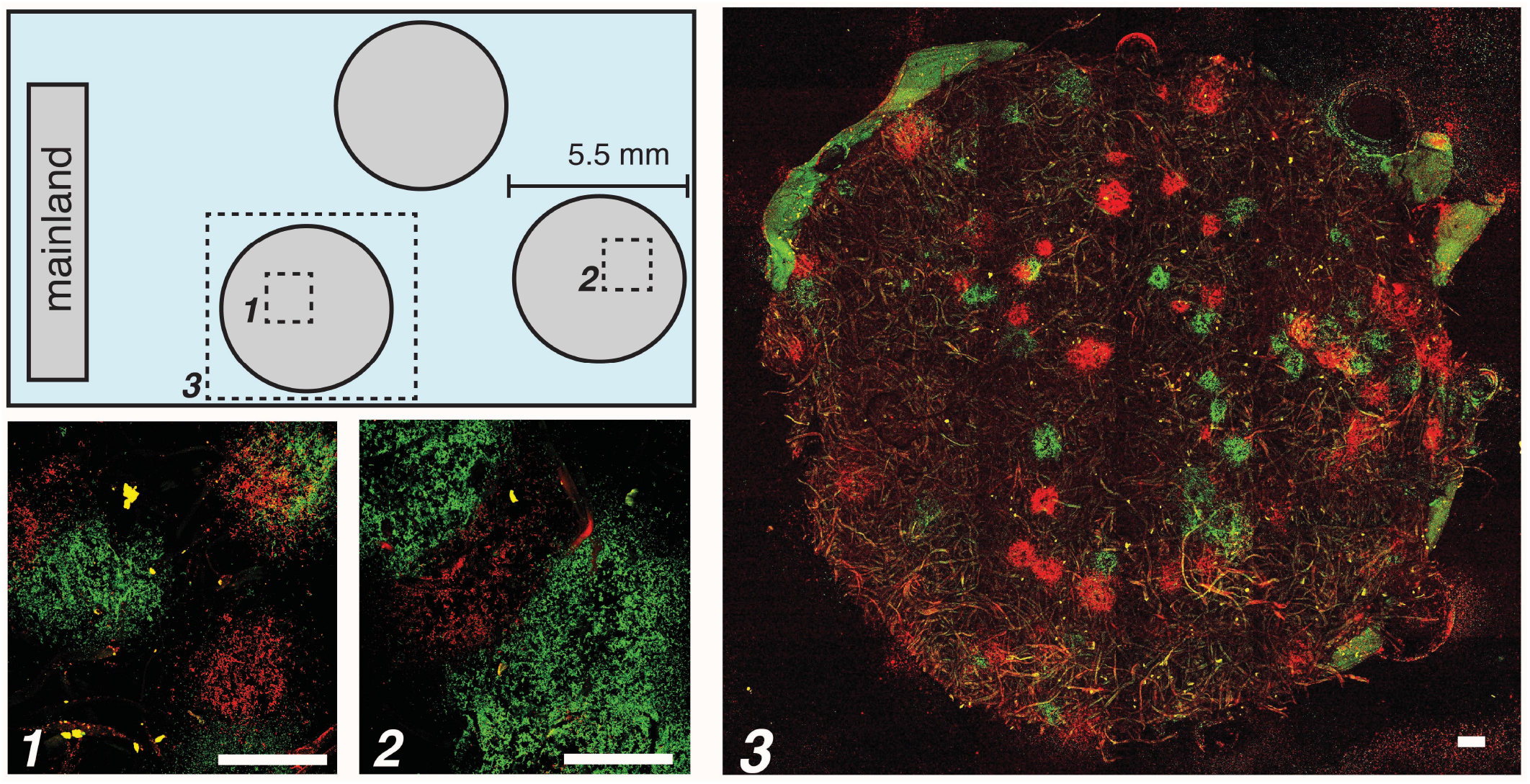
Colonization of an archipelago. *E. coli* inoculated on the mainland colonize an initially uninhabited archipelago of three paper islands. Paper scaffolds are surrounded by liquid minimal medium, and (non-inoculated) islands were pre-treated with glucose which starts to diffuse out of the scaffolds upon wetting. Diffusing glucose promotes bacterial migration by creating a temporary glucose gradient increasing towards the non-inoculated islands. Confocal scans correspond to the areas indicated with 1, 2, and 3 in the cartoon. Scale bars are 200 *µ*m.

Branching networks and archipelagos are canonical landscapes in ecology. Using no more than paper and scissors, such diverse ecological scenarios can now be explored in habitats structured at the microscopic scales relevant to bacterial ecology. Given the ease with which paper can be cut in milli-to centimeter shapes, the platform presented here can be used to address a wide range of questions on how multi-scale landscape geometry and topology affect bacterial community dynamics. In addition to spatial structure, resource heterogeneity can be incorporated by seeding nutrients locally in the paper giving rise to a rich repertoire of ecosystems that can be modeled in paper. By providing a versatile, easy-to-use, and virtually zero-cost alternative to microfabrication-based approaches to experimental microbial ecology, this work fits in a broader push towards democratizing science by eliminating the need for expensive and specialized equipment by providing inexpensive alternatives that rely on generic materials and tools.

## MATERIALS & METHODS

### Preparation and inoculation of paper scaffolds

Landscape geometry was designed in Adobe Illustrator CS6 and cut in Whatman 1 Chr chromatography paper (0.18 mm thick) using a laser cutter (Versa Laser-Universal Laser VL-300). Cut paper was autoclaved and dried before use.

#### Branching assay

Separate overnight cultures of *E. coli* strain JEK1036 (W3110 lacYZ::GFPmut2) and strain JEK1037 (W3110 lacYZ::mRFP) were diluted 1/200 in fresh LB medium supplemented with 10 *µ*M isopropyl *β*-D-1-thiogalactopyranoside (LB-IPTG), grown to mid log phase, and mixed at 1:1 ratio for inoculation (pre-mixing density measured by optical density). Paper scaffolds were submerged in LB-IPTG for 5 minutes, excess medium was allowed to drip from the paper, and the medium saturated paper was suspended horizontally on a grille of thin peek tubing (5 mm pitch). The assembly was transferred to a chamber with saturated humidity and placed in an incubator set to 37*°*C for 30 minutes prior to inoculation. The 1:1 mix of GFP and RFP labelled *E. coli* was inoculated onto the center of the scaffold using a 1 microliter inoculation loop giving rise to an inoculation zone of approximately 2 millimeters. Scaffolds were incubated for 15 hours at 37*°*C.

#### Archipelago assay

Separate overnight cultures of *E. coli* strain JEK1036 (W3110 lacYZ::GFPmut2) and strain JEK1037 (W3110 lacYZ::mRFP) were diluted 1/200 in fresh M9 medium supplemented with 0.4% glucose and 10 *µ*M IPTG and grown to mid log phase. Mid log phase cells were washed by spinning down, discarding the supernatant, and re-suspending in M9-IPTG without glucose, washed cells were mixed at 1:1 ratio for inoculation. The paper scaffolds comprising the archipelago (1 mainland, 3 islands) were positioned on a layer of parafilm on top of a cover slide. The parafilm was cut around the paper, and excess parafilm (i.e. parafilm not sandwiched between paper and glass) was removed. The slide was heated on a hot-plate to briefly melt the parafilm and secure the scaffolds to the glass slide. After cooling 5 *µ*L of a 20% glucose solution was pipetted onto the paper islands (not the mainland) and let to dry. The 1:1 mix of GFP and RFP labelled *E. coli* was inoculated onto the mainland using a 1 microliter inoculation loop. A rectangular Gene Frame adhesive (17 x 28 x 0.25 mm, Thermo Fisher Scientific) was placed around the paper scaffolds, and 125 *µ*L of M9 medium with 10 *µ*M IPTG (no glucose) was introduced. The archipelago was closed by placing a cover slip on the Gene Frame and incubated at 37*°*C for 15 hours.

### Confocal imaging of bacteria in paper

Archipelagoes were imaged without modification. Branching scaffolds were fixed by submerging in 4% formaldehyde in phosphate buffered saline (PBS) for 15 minutes and subsequently washed 3 times with PBS. Scaffolds were imaged in PBS + 50% glycerol in an imaging chamber (cover slide, Gene Frame, cover slip). Imaging was performed on a Nikon A1R Confocal system controlled using NIS-Elements C software at 10x or 20x magnification.

### qPCR of bacteria in paper

Fragments (2 mm^2^) were cut from the extremities of the branching and non-branching landscapes using a razor blade and used for genomic DNA extraction (NucleoSpin Tissue kit, Macherey-Nagel). Quantitative PCR was performed in duplicate on an Eco Real-Time PCR system (Illumina) using SYBR Green PCR Master Mix (ThermoFisher Scientific) and primer sets GFP: 5’ GGCACTCTTGAAAAAGTCATGCT - 3’ (forward), 5’ - CCATGGCCAACACTTGTCACT - 3’ (reverse), RFP: 5’ - CCCTGAAGGGCGAGATCAA - 3’ (forward), 5’ TGGCCATGTAGGTGGTCTTG - 3’ (reverse). Data was analyzed in MATLAB 2016a using a custom script.

## SUPPLEMENTARY INFORMATION

**Supplementary Movie 1.** Movie of a confocal Z-stack starting from the paper surface moving 33.6 micrometer into the paper in 4.8 micrometer increments. Green/red mixed cellular aggregates can be seen through the paper scaffold.

## ACKNOWLEDGEMENTS

We thank Bobak Mosadegh, Karen Simon, Jonathan Hennek, and Gulden Camci-Unal (Whitesides Lab) and Fabai Wu, Jakub Wiktor, and Jaco van der Torre (Dekker Lab) for fruitful discussion. FJHH acknowledges support from a Kavli Exchange Fellowship, and a Burroughs Wellcome Fund Career Award at the Scientific Interface. CD acknowledges support from ERC Advanced Grants SynDiv (No. 669598) and Nanoforbio (No. 247072), and the Netherlands Organization of Scientific Research (NWO/OCW) as part of the Frontiers of Nanoscience Program.

